# Proteome Dynamics Across the Blastogenic Cycle of *Botryllus schlosseri* Reveals Targets for Cell Immortalization

**DOI:** 10.1101/2025.07.01.662660

**Authors:** Weizhen Dong, Maxime Leprêtre, Isabel Enriquez, Brenda Luu, Mandy Lin, Jens Hamar, Dietmar Kültz

**Affiliations:** Department of Animal Sciences and Genome Center, University of California - Davis, One Shields Ave., Meyer Hall, Davis, CA, 95616, USA

**Keywords:** mass spectrometry, proteomics, *Botryllus schlosseri*, cell proliferation, asexual reproduction, tunicates

## Abstract

The colonial tunicate *Botryllus schlosseri* regenerates weekly through a cyclical process in which adult zooids are replaced by a new generation of buds. While this dynamic asexual development is a hallmark of the species, its molecular regulation remains poorly understood. This study presents the first comprehensive proteomic analysis of *B. schlosseri* blastogenesis at the individual zooid level, using data-independent acquisition mass spectrometry to quantify protein abundance across developmental stages. The results reveal extensive proteome remodeling between proliferating buds and degenerating zooids. Co-expression analysis identified stage-specific protein modules enriched for biosynthesis and cell cycle pathways in buds, and for apoptosis, catabolism, and metabolic remodeling in zooids. A focused comparison between takeover buds and takeover zooids uncovered distinct regulatory programs controlling proliferation and senescence. Key proteins, including CDK1, CDK2, HDAC2, and PCNA, were identified as candidate regulators of cell cycle progression. These findings provide a molecular framework for understanding regeneration in a basal chordate and offer protein targets that may enable cell cycle re-entry and long-term culture of tunicate primary cells.

**Summary Statement:** This study maps proteome dynamics during the blastogenic cycle in *Botryllus schlosseri*, identifying candidate proteins that regulate cell proliferation and offer targets for tunicate cell line development.

## Introduction

The colonial tunicate *Botryllus schlosseri* (Fig. 1A) has emerged as a compelling model organism for exploring the mechanisms of regeneration (Voskoboynik *et al*., 2007; Ricci *et al*., 2022), aging (Munday *et al*., 2015; Voskoboynik and Weissman, 2015), and stress resilience (Dijkstra and Simkanin, 2016; Tasselli *et al*., 2017). As the closest living invertebrate taxon relative to vertebrates (Delsuc *et al*., 2006), tunicates occupy a critical phylogenetic position that bridges evolutionary milestones, offering unique insights into the molecular underpinnings responsible for conservation, innovation, and loss of cellular and organismal processes during chordate phylogeny (Fig. S1). Botryllid tunicates stand out as the chordate phylogenetically closest to humans that are capable of whole-body regeneration during asexual reproduction and in response to injury, a trait that has been lost in all vertebrates and most other chordates. These characteristics render *B. schlosseri* a unique model for studying molecular mechanisms that promote tissue regeneration, cell proliferation, and cell differentiation in chordates.

**Figure 1.**
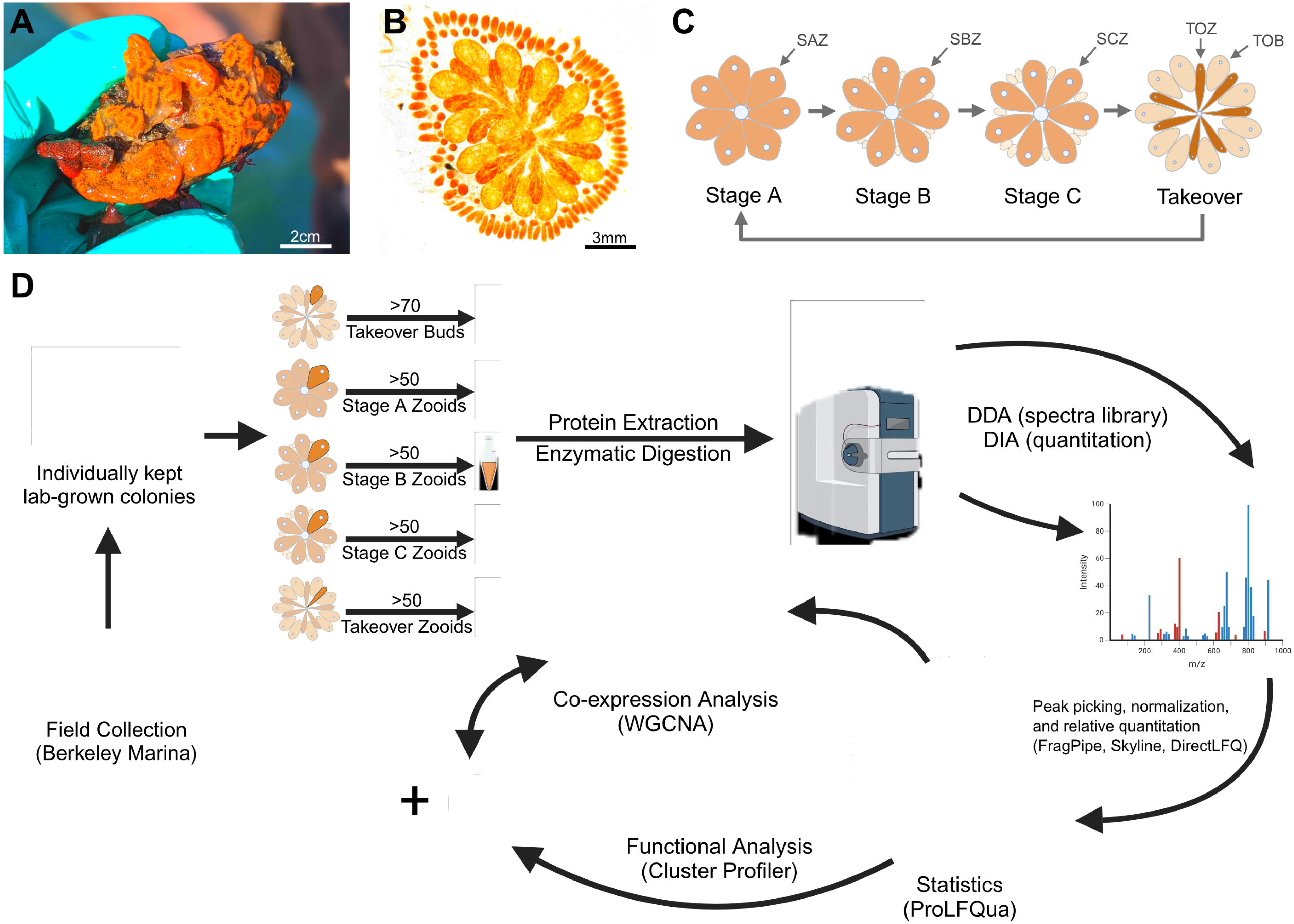
Overview of the blastogenic cycle and proteomics workflow in *Botryllus schlosseri*. (A) Photograph of *B. schlosseri* colonies collected from the Berkeley Marina Harbor, growing on a mussel shell. Individual zooids within each colony measure up to 3 mm in length. (B) Microscopic view of a *B. schlosseri* colony during the takeover stage. Senescing takeover zooids (TOZ) appear darker reddish than the proliferating takeover buds (TOB), which appear light orange. (C) Schematic illustration of the blastogenic cycle of *B. schlosseri*, highlighting the key stages at the individual animal level: Stage A Zooid (SAZ), Stage B Zooid (SBZ), Stage C Zooid (SCZ), Takeover Zooid (TOZ), and Takeover Bud (TOB). (D) Workflow for proteomic analysis of *B. schlosseri* colonies. Colonies are collected from the field, maintained individually in laboratory conditions, and sampled at specific blastogenic stages. Samples undergo protein extraction, enzymatic digestion, and mass spectrometry analysis using both Data-Dependent Acquisition (DDA) for spectral library generation and Data-Independent Acquisition (DIA) for quantitation. Data normalization (DirectLFQ), statistical analysis (ProLFQua), and co-expression analysis (WGCNA) are performed, followed by functional enrichment using KEGG and Gene Ontology (GO) tools. Created in BioRender. Dong, V. (2025) https://BioRender.com/q48egyl.

Notably, *B. schlosseri* undergoes a synchronized weekly blastogenic cycle of asexual reproduction, during which old zooids degenerate and are replaced by new primary buds. First described by (Sabbadin, 1955), and then later reclassified and characterized in detail by (Manni *et al*., 2007), this cycle consists of four stages (A-D) and occurs within a colony where three generations coexist, connected by a communal vasculature. Adult zooids actively feed, while primary and secondary buds remain nutritionally dependent on the zooids. As the cycle progresses, primary buds develop through stages A to C, culminating in the takeover stage D (Fig. 1B), where synchronized zooid regression and increased cellular turnover drive colony renewal. This cyclical process mirrors multiple aspects of growth, maturation, and programmed cell death common to vertebrates (Anselmi *et al*., 2022), thus providing an *in vivo* model for understanding conserved mechanisms of cellular turnover in chordates. Moreover, studying blastogenic takeover and injury-induced whole-body regeneration in botryllid tunicates offers an opportunity to identify molecular processes of chordates that can be targeted to promote tissue and whole-body regeneration by genetic or pharmacological intervention in other chordates that have lost the ability of comprehensive tissue and whole-body regeneration during evolution.

However, key limitations have impeded deeper mechanistic studies of *B. schlosseri*, including the scarcity of information on the proteome (Kültz *et al*., 2024), which represents the main determinant of cellular and organismal structure and function, and the absence of a established cell line models. Addressing these gaps is essential for advancing functional investigations and for high-throughput genetic manipulation of cellular processes in a controlled environment to establish causality between gene function and phenotype. Despite its utility as an *in vivo* model, the lack of *B. schlosseri* cell lines limit the ability to manipulate and investigate cellular mechanisms in a controlled environment by high-throughput genetic engineering and other causality-aimed approaches. While some species and cell types can undergo spontaneous immortalization without the introduction of foreign elements (Vaughn *et al*., 1977; Saad *et al*., 2023), most species are believed to require targeted interventions to achieve stable, long-term cell growth. With current approaches, isolated *B. schlosseri* primary cells exit the G1 phase of the cell cycle and remain quiescent in G0 after relatively short-term (1-3 weeks) primary culture and optimal conditions for long-term culture remain undefined (Rabinowitz and Rinkevich, 2004; Qarri *et al*., 2023).

Overcoming these challenges requires innovative strategies to enhance cell survival and proliferation while inhibiting senescence *in vitro*. Mammalian cell lines have been successfully immortalized using viral oncoproteins, such as the SV40 large T antigen, or modifications of the cell cycle machinery, including the introduction of human telomerase reverse transcriptase (hTERT) (Linzer and Levine, 1979; Hawley Nelson *et al*., 1989; Bodnar *et al*., 1998). However, such methods often fail in non-mammalian systems, where conserved pathways like cell cycle regulation may rely on subtly divergent regulatory mechanisms that are still poorly characterized in aquatic invertebrates. To date, the only established strictly marine invertebrate cell lines are a sponge-derived cell line derived from the phylum Porifera by spontaneous immortalization (Hesp *et al*., 2023), and a hybrid shrimp cell line, PmLyO-Sf9, created by fusing *Penaeus monodon* lymphoid cells with Sf9 insect cells (Anoop *et al*., 2021).

Terrestrial invertebrate models have offered limited insight. *Drosophila* cells, for example, could be immortalized by overexpressing Ras^V12^, but not Myc, highlighting that even targeting well-established oncogenes has varying effectiveness across species^21^. These outcomes underscore the importance of identifying species-specific targets rather than exclusively extrapolating target identification based on knowledge of mammalian or terrestrial invertebrate systems. Even core regulators such as cyclin-dependent kinases (CDKs) differ across lineages and primitive chordates have fewer copies than mammals due to CDK gene duplication events during evolution. For instance, yeast encodes only a single CDK (Cdc28) (Nasmyth, 1993), while humans possess 21 CDK paralogs that include CDK4, a common target in mammalian immortalization protocols(Chotiner, Wolgemuth and Wang, 2019). Moreover, CDK1 gene duplication has been reported in the tunicate *Oikopleura* (Ma, Øvrebø and Thompson, 2022). Such variability suggests that successful strategies in non-mammalian systems will require a deeper understanding of their unique regulatory landscapes. Identification of species-specific molecular proteome signatures associated with stages of active proliferation and senescence will aid in the identification of potent regulators of cell growth and survival.

Previous transcriptomic studies in *B. schlosseri* have provided valuable insight into pathways involved in stem cell activation, immune responses, and oxidative stress responses associated with aging(Ben-Hamo *et al*., 2018; Rosental *et al*., 2018; Goldstein *et al*., 2021; Rodriguez *et al*., 2021). Apoptosis and autophagy have also been implicated in the degeneration of adult zooids, supporting the tissue turnover required for bud development (Cima *et al*., 2010; Franchi *et al*., 2016). However, these studies did not directly capture the proteins responsible for proliferation and senescence phenotypes during the blastogenic cycle. Considering the proteome is necessary due to imperfect correlation of mRNA abundance and protein abundance regulation in mammalian and other chordate cells (Schwanhäusser *et al*., 2011; Buccitelli and Selbach, 2020; Wang *et al*., 2020; Root *et al*., 2021). Furthermore, transcriptomic analyses do not capture functional protein-level activity and post-translational modifications, which are critical for understanding dynamic changes in cellular phenotypes. Proteomic analyses, on the other hand, enable direct investigation of protein abundances, interactions, and modifications, offering a more comprehensive and functional understanding of cellular responses. Recent comparative proteomic studies have demonstrated the superior ability of proteomics to elucidate complex stress responses, such as salinity adaptation, by capturing dynamic protein-level adjustments that transcriptomics approaches alone would miss (Leprêtre *et al*., 2025). Recent advances in mass spectrometry allow the quantification of thousands of proteins in a single sample, making it possible to analyze whole proteome changes across different experimental conditions. This enables the discovery and identification of new molecular targets that promote phenotypes of interest.

By leveraging *B. schlosseri*’s unique biology and label-free quantification DIA proteomics, this study aims to comprehensively characterize proteome dynamics across different blastogenic stages to identify key proteins and molecular signatures associated with *B. schlosseri* cell proliferation and senescence. Initially, proteomics was performed across all blastogenic stages to capture global protein abundance regulation and identify corresponding functional adjustments during blastogenesis. Subsequently, a more focused proteomic follow-up study was conducted focusing on the blastogenic cycle stages that are most informative regarding critical regulators of cell proliferation and senescence.

## Results

### Distinct Proteomic Landscapes Throughout the Blastogenic Cycle

Adult zooids and primary buds at specific blastogenic stages were classified as follows: Stage A zooids (SAZ), Stage B zooids (SBZ), Stage C zooids (SCZ), Takeover zooids (TOZ), and Takeover buds (TOB) (Fig. 1C). Secondary buds, due to their extremely small size, could not be physically separated from the primary buds and are assumed to contribute minimally to the primary bud data. Additionally, primary buds from stages A, B, and C were excluded due to the challenge of isolating sufficient protein for analysis.

To establish a foundational proteomics analysis of *Botryllus schlosseri* at the individual animal zooid level, the proteomes of adult zooids across four blastogenic stages (SAZ, SBZ, SCZ, and TOZ) and emerging buds from the takeover stage (TOB) were examined. To ensure that observed differences reflect biological variation rather than clonal effects, all colonies were confirmed to represent unique genotypes prior to sample collection (Fig. S2). The experimental workflow, including sample preparation and proteomics analysis, is outlined in Fig. 1D. A total of 15,156 peptides and 3,155 proteins were reliably quantified across all stages. Among the identified proteins, 45% (1,432) exhibited statistically significant changes in abundance across stages (ANOVA, FDR < 0.1), indicating extensive proteomic remodeling throughout the blastogenic cycle.

Principal Component Analysis (PCA) performed on these differentially abundant proteins (DAPs) demonstrated clear separation of expression patterns among blastogenic stages, with TOB exhibiting the most distinct proteome compared to adult zooids at all other stages (Fig. 2A). PC1, which explains 64% of the total variance, clearly discriminates TOB, SAZ, and TOZ, reflecting the major proteomic shifts associated with the transition from active adult zooids to degenerating zooids and the emergence of new primary buds. PC2 captures more subtle differences between SBZ and SCZ, corresponding to the progressive maturation of adult zooids prior to takeover. The distinct separation observed in PCA underscores the molecular coordination driving developmental renewal and programmed cell death in *B. schlosseri*.

**Figure 2.**
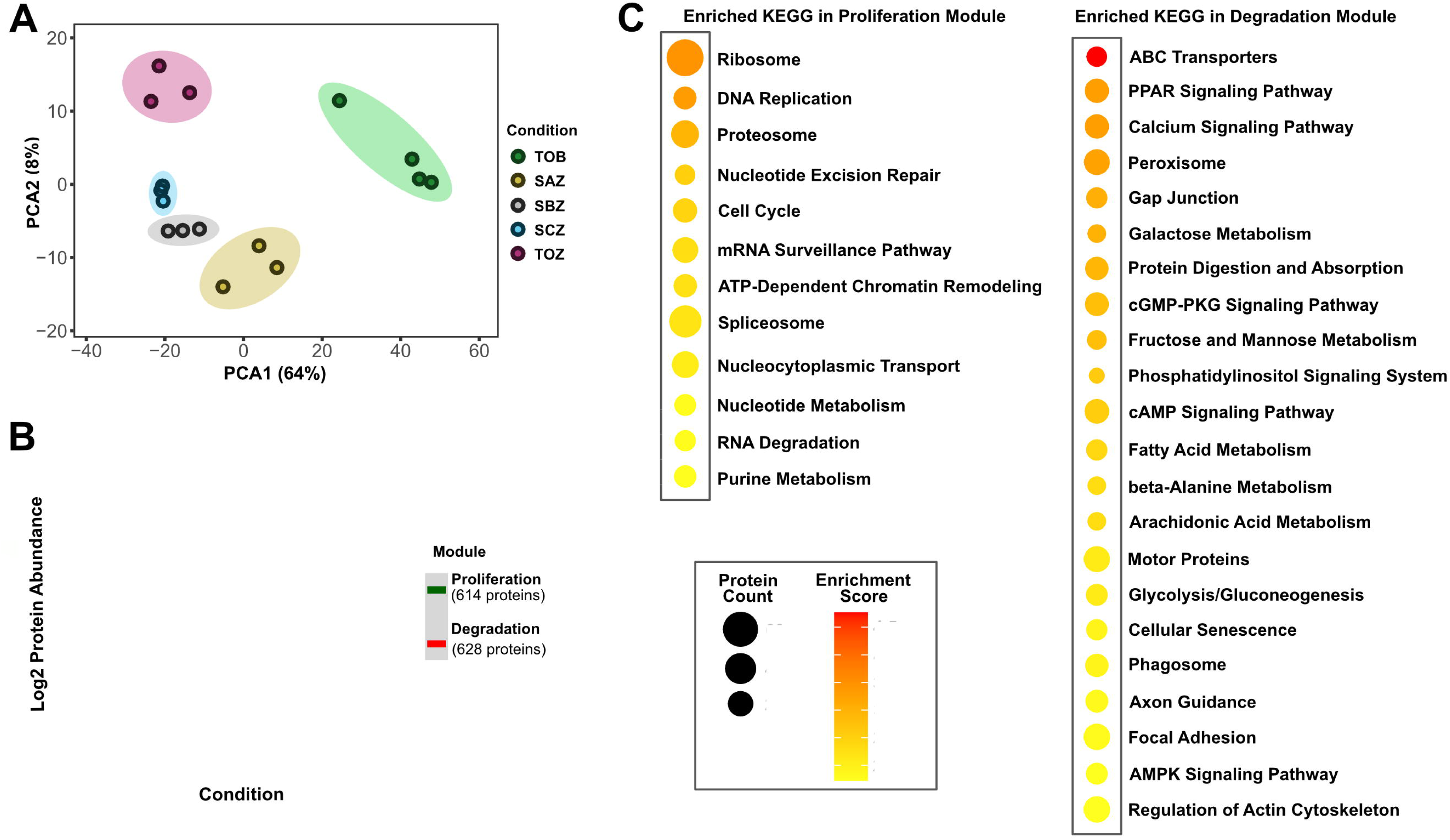
Global proteomics analysis of *Botryllus schlosseri* across blastogenic stages (Study 1). (A) Principal component analysis (PCA) of five blastogenic stages, TOB (Takeover Bud), SAZ (Stage A Zooid), SBZ (Stage B Zooid), SCZ (Stage C Zooid), and TOZ (Takeover Zooid), based on 1,432 proteins significantly differentially expressed by ANOVA (FDR < 0.1). (B) Weighted Gene Co-expression Network Analysis (WGCNA) identified two major co-expression modules across blastogenic stages: a Proliferation module (green, 614 proteins) and a Degradation module (red, 628 proteins). The plot shows the average logl_ protein abundance for each module across stages, with shaded areas representing standard deviation (n = 3). (C) KEGG pathway enrichment results for proteins in the Proliferation (left) and Degradation (right) modules. Circle size represents the number of proteins mapped to each pathway, while color reflects enrichment score. Pathways specific to vertebrate physiology, human disease, or dependent on cell types absent in *B. schlosseri* were excluded prior to visualization.

### Co-expression Modules Reveal Distinct Functional Programs

Weighted Gene Co-expression Network Analysis (WGCNA) performed on the DAPs identified two major modules with distinct expression trends across the blastogenic cycle (Fig. 2B). The first module, shown in green and termed the proliferation module, contains 614 proteins whose abundance decreases from the TOB stage to the TOZ stage. In contrast, the second module, shown in red and referred to as the degradation module, includes 628 proteins that increase in abundance over the same transition. These opposing trends highlight a coordinated switch from proliferative to degradative cellular programs as the cycle progresses. The remaining 190 DAPs were not classified into any co-expression module. Notably, protein abundance shifted most dramatically between the TOB and SAZ stages, with DAPs exhibiting modulations averaging a log fold change (FC) of 1, whereas subsequent transitions were more gradual.

To interpret the biological relevance of these co-expression patterns, Kyoto Encyclopedia of Genes and Genomes (KEGG) pathway enrichment analysis was performed for each module (Fig. 2C). Pathways associated with vertebrate-specific physiology, human diseases, or cell types absent in *B. schlosseri* were excluded from this analysis. The proliferation module (Fig. 2B, green, 614 proteins), which decreases in abundance from TOB to TOZ, was broadly enriched for pathways related to cellular proliferation, such as translational capacity, cell cycle progression, and macromolecular biosynthesis. The most enriched pathways were ribosome biogenesis (70 DAPs), DNA replication (14 DAPs) and proteasome function (27 DAPs). Other enriched pathways included groups involved in cell cycle progression, macromolecular biosynthesis, mRNA surveillance, chromatin remodeling, and nucleocytoplasmic transport, reflecting coordinated regulation of processes essential for cell growth and division. In contrast, the degradation module (Fig. 2B, red, 628 proteins), which increases from TOB to TOZ, was enriched for catabolic and metabolic remodeling pathways. The most significant enrichment was seen in ABC transporters (10 DAPs), followed by PPAR signaling (17 DAPs), calcium signaling (17 DAPs), peroxisome (21 DAPs), gap junction (11 DAPs), Galactose metabolism (8 DAPs) and protein digestion and absorption (15 DAPs). These pathways likely support nutrient salvage, stress signaling, and tissue breakdown during zooid regression.

### Confirmation of Pro-and Anti-Proliferative Protein Modules During Takeover

To support the findings from the first study, a second study was conducted, focusing on the three most biologically distinct stages identified earlier: the onset of proliferation (TOB), the adult stage (SAZ), and peak regression (TOZ). To increase statistical power and improve proteome coverage, the number of biological replicates was expanded from three to seven per condition. This second study yielded 22,458 peptides and 4,015 proteins, representing a 32.5% increase in peptide detection and a 27% increase in protein identification compared to the first study. Of the detected proteins, 2,919 were shared across both studies, accounting for 92.5% of the first and 72.7% of the second study. An additional 1,096 proteins were uniquely identified in the second study, while 236 were exclusive to the first (Fig. 3A). The improved detection is likely due to both the increased number of replicates and the use of a more inclusive spectral library, which incorporated both DIA and MS1 (Pino *et al*., 2020), in contrast to the DDA-based approach used previously. PCA of the 2,020 DAPs identified from the second study showed clear separation among TOB, SAZ, and TOZ, consistent with the pattern observed in the first study, with TOB and TOZ exhibiting the most distinct proteome differences (Fig. 3B).

**Figure 3.**
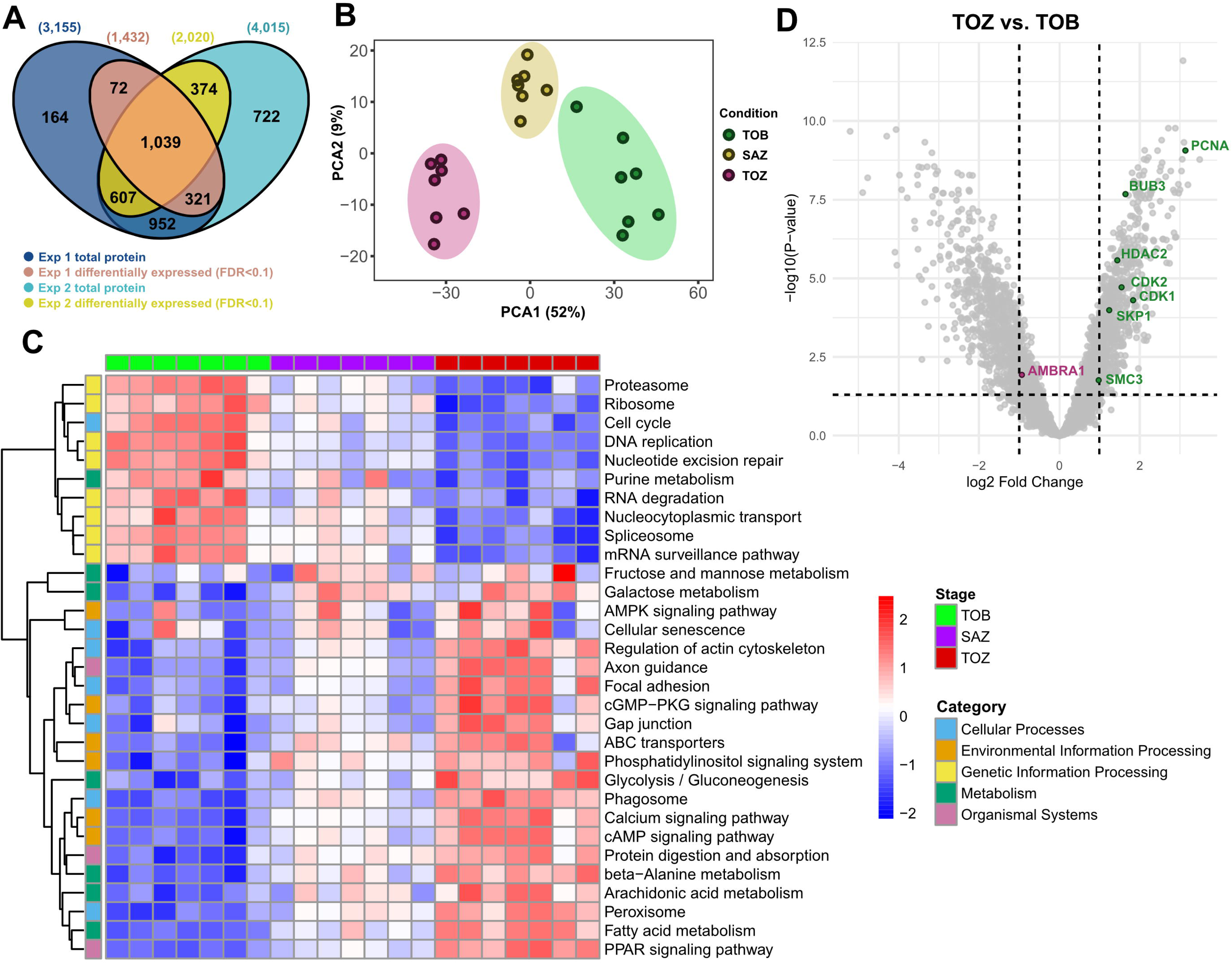
Global proteomics analysis of *Botryllus schlosseri* across blastogenic stages (Study 2). (A) Venn diagram showing the overlap of total and differentially expressed proteins (FDR < 0.1) between two independent experiments. (B) Principal component analysis (PCA) based on 2,020 proteins significantly differentially expressed by ANOVA (FDR < 0.1), showing clear separation of samples from three blastogenic stages: TOB (Takeover Bud), SAZ (Stage A Zooid), and TOZ (Takeover Zooid). (C) Heatmap displaying row-scaled logl_-summed protein abundances for annotated KEGG pathways (rows) across biological replicates of TOB, SAZ, and TOZ (columns). Color intensity reflects Z-scores, with red indicating higher and blue indicating lower relative abundance within each pathway. Top annotation bars indicate sample stage, while left-side color bars categorize pathways by KEGG functional category. Hierarchical clustering of rows and columns was performed using Euclidean distance and complete linkage. Samples are ordered by stage to highlight temporal dynamics in pathway activity. (D) Volcano plot showing differential protein expression between TOB and TOZ. Each dot represents a single protein, plotted by logl_ fold change (x-axis) and –logl_l_(FDR) (y-axis). Vertical dashed lines indicate logl_ fold change thresholds of ±1; the horizontal dashed line indicates an FDR threshold of 0.05. Labeled proteins highlight candidates with established or putative roles in cell cycle progression, biosynthesis, and remodeling, including CDK1, CDK2, PCNA, HDAC2, BUB3, SKP1, SMC3, and AMBRA1.

To assess whether the pathway-level trends identified in the first study were conserved, the summed log protein abundance for each KEGG pathway enriched in the first study was calculated in the second study (n = 7) and visualized across TOB, SAZ, and TOZ (Fig. 3C). This analysis revealed distinct stage-specific patterns in pathway activity, which grouped into two major expression clusters. Pathways associated with cellular proliferation, including proteasome function, ribosome biogenesis, cell cycle regulation, DNA replication, nucleotide excision repair, purine metabolism, RNA degradation, and the spliceosome, showed the highest abundance in TOB, followed by reduced levels in SAZ and further decreases in TOZ. In contrast, a second group of pathways, including AMPK signaling, cellular senescence, actin cytoskeleton regulation, ABC transporters, calcium signaling, protein digestion and absorption, and fatty acid metabolism, displayed the highest abundance in TOZ, consistent with their roles in tissue breakdown and metabolic recycling during zooid regression. The heatmap also included all seven replicates per stage, revealing strong clustering by stage and indicating minimal genotypic variation among individuals in this study.

### Cell Cycle–Associated Proteins

To identify proteins that may regulate the transition between proliferation and senescence, stage-specific expression was compared between TOB and TOZ, which are the two stages that showed the greatest differences in protein abundance. DAPs that contributed to the cell cycle KEGG pathway (ko04110) were further examined. To reduce redundancy, isoforms that shared the same KEGG annotation were collapsed, retaining a single representative per protein. Using a significance threshold of FDR < 0.05 for TOB versus TOZ comparisons, 16 DAPs were identified to be associated with the cell cycle KEGG pathway (Table 1). All the 16 DAPs showed increased abundance in TOB compared to TOZ and are highlighted in the volcano plot shown in Fig. 3D. These included proteins involved in DNA replication, cell cycle progression, checkpoint control, chromosomal cohesion, chromatin remodeling, and the 14-3-3 family (Fig. 4).

**Figure 4.**
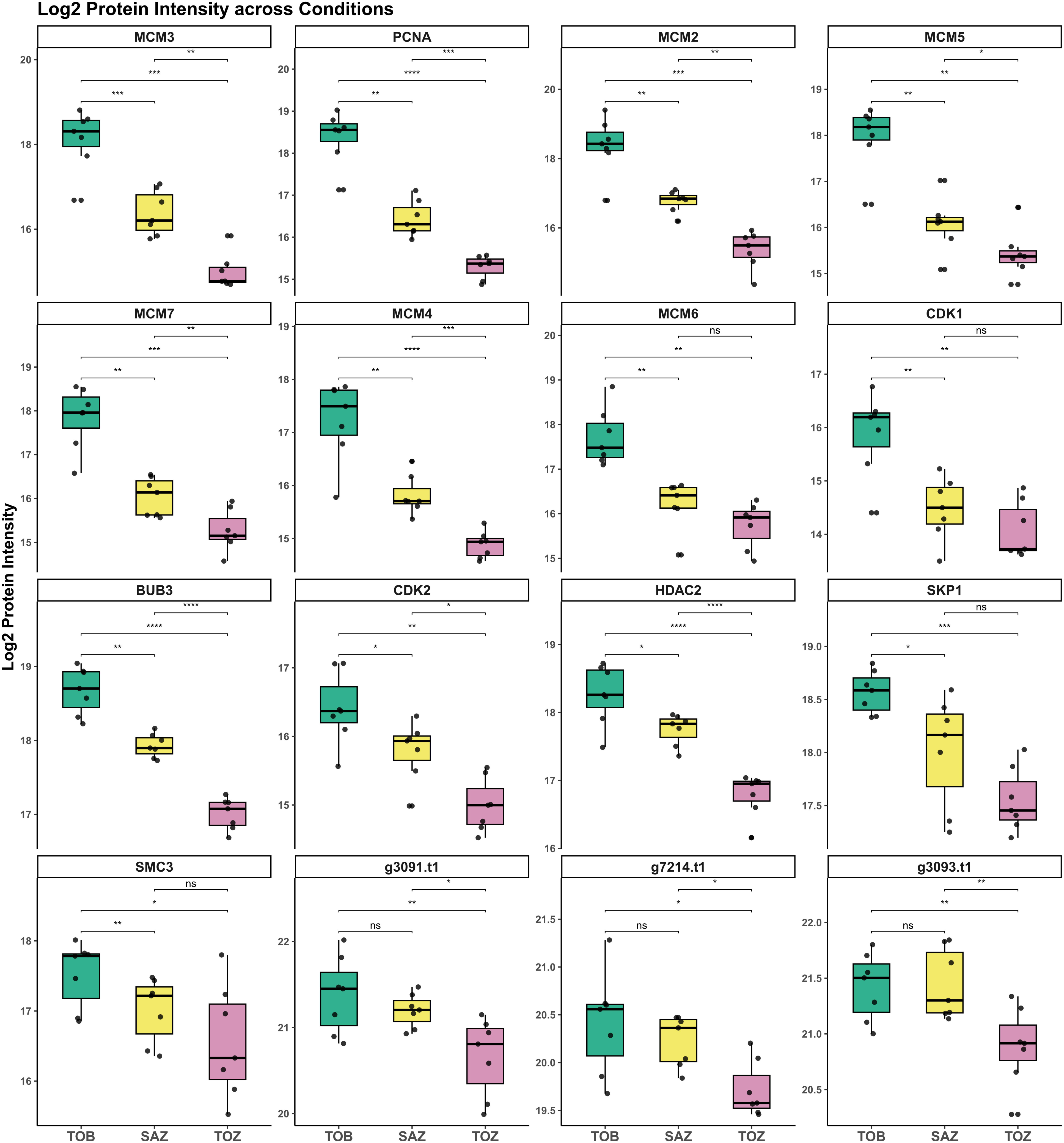
Targeted expression analysis of cell cycle–associated proteins during key stages of the *B. schlosseri* blastogenic cycle. Boxplots display logl_-transformed protein intensities for selected regulators across the Takeover Bud (TOB), Stage A Zooid (SAZ), and Takeover Zooid (TOZ) stages. Proteins include core components of DNA replication (MCM2–7 complex, PCNA), cell cycle progression (CDK1, CDK2, SKP1), chromosomal cohesion and checkpoint control (SMC3, BUB3), chromatin remodeling (HDAC2), and three 14-3-3 homologues.

**Table 1.**
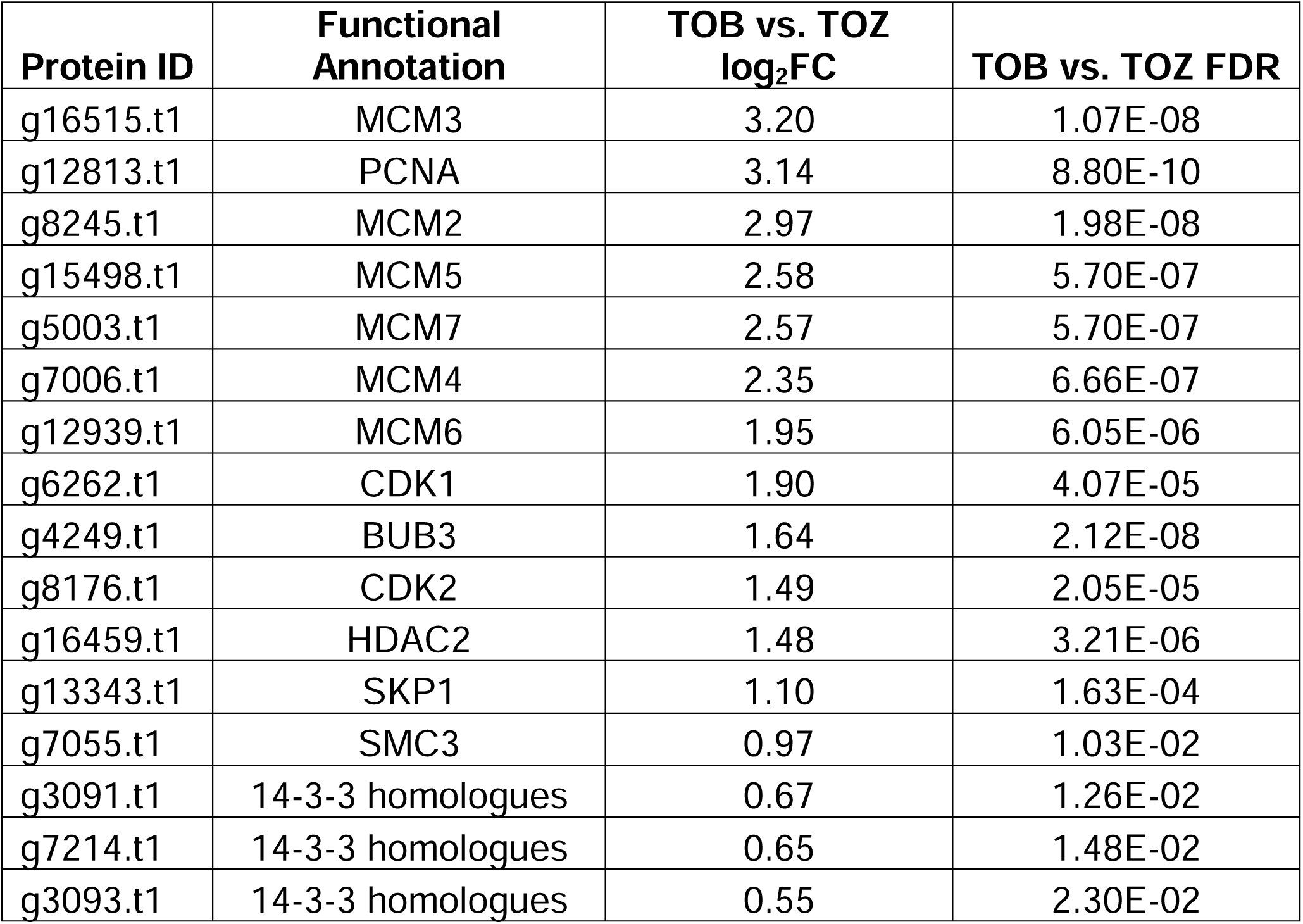
Proteins annotated to the cell cycle KEGG pathway that are significantly enriched in takeover buds (TOB) relative to takeover zooids (TOZ). Abbreviations: logl_FC, logl_-transformed fold change; FDR, false discovery rate. Proteins were considered significantly enriched with FDR < 0.05.

| Protein ID | Functional Annotation | TOB vs. TOZ $\log_2\text{FC}$ | TOB vs. TOZ FDR |
| --- | --- | --- | --- |
| g16515.t1 | MCM3 | 3.20 | 1.07E-08 |
| g12813.t1 | PCNA | 3.14 | 8.80E-10 |
| g8245.t1 | MCM2 | 2.97 | 1.98E-08 |
| g15498.t1 | MCM5 | 2.58 | 5.70E-07 |
| g5003.t1 | MCM7 | 2.57 | 5.70E-07 |
| g7006.t1 | MCM4 | 2.35 | 6.66E-07 |
| g12939.t1 | MCM6 | 1.95 | 6.05E-06 |
| g6262.t1 | CDK1 | 1.90 | 4.07E-05 |
| g4249.t1 | BUB3 | 1.64 | 2.12E-08 |
| g8176.t1 | CDK2 | 1.49 | 2.05E-05 |
| g16459.t1 | HDAC2 | 1.48 | 3.21E-06 |
| g13343.t1 | SKP1 | 1.10 | 1.63E-04 |
| g7055.t1 | SMC3 | 0.97 | 1.03E-02 |
| g3091.t1 | 14-3-3 homologues | 0.67 | 1.26E-02 |
| g7214.t1 | 14-3-3 homologues | 0.65 | 1.48E-02 |
| g3093.t1 | 14-3-3 homologues | 0.55 | 2.30E-02 |

DNA replication factors exhibited the most pronounced changes with proliferating cell nuclear antigen (PCNA) showed one of the greatest differential abundance between TOB and TOZ (log FC = 3.1, FDR = 8.8 × 10 ¹). All six subunits of the minichromosome maintenance complex (MCM2, MCM3, MCM4, MCM5, MCM6, and MCM7), which form the core replicative helicase required for DNA unwinding during S phase, were greatly upregulated in TOB, suggesting increased licensing and replication initiation activity. Cyclin-dependent kinase 1 (CDK1) and cyclin-dependent kinase 2 (CDK2) were also significantly upregulated (log FC = 1.9 and 1.5; FDR = 4 × 10 and 2 × 10), consistent with progression through the G2/M and G1/S phases, respectively. S-phase kinase-associated protein 1 (SKP1), a core component of the SCF E3 ubiquitin ligase complex, was increased in abundance in TOB (log FC = 1.1, FDR = 1.6 × 10). Proteins involved in mitotic checkpoint control and chromatid cohesion were also enriched. Structural maintenance of chromosomes protein 3 (SMC3), a subunit of the cohesin complex necessary for sister chromatid pairing, was more abundant in TOB (log FC = 1.0, FDR = 0.01). BUB3 mitotic checkpoint protein (BUB3), a conserved spindle assembly checkpoint regulator, showed high regulation in TOB (log FC = 1.6, FDR = 2.1 × 10), reflecting the need for mitotic surveillance during rapid proliferation. Histone deacetylase 2 (HDAC2), a class I histone modifier involved in chromatin compaction, was also significantly upregulated in TOB (log FC = 1.5, FDR = 3.2 × 10). Notably, HDAC1 was not detected in the study, suggesting a potentially dominant role for HDAC2 in regulating chromatin state during cell cycle transitions in this system. Finally, several proteins annotated as members of the 14-3-3 protein family showed increased abundance in TOB. Although their specific isoforms could not be resolved from the blastp annotation, their enrichment suggests a potential regulatory role in checkpoint signaling or cell cycle coordination during blastogenesis.

## Discussion

### Proteome Changes During the Blastogenic Cycle

The extensive proteomic remodeling which composed of nearly half of the detected proteome observed across the blastogenic cycle of *Botryllus schlosseri* reflects major biological transitions that define its asexual reproduction. PCA results of the first study showed that the while each of the four adult stage are distinct from each other in terms of their proteome, the most significant molecular changes happen at the takeover stage between TOZ and TOB where adult zooids degenerate and new buds develop. Co-expression analysis also corroborate with this pattern, the sharp co-expression transition from TOB to SAZ from both modules reflects the accelerated progression of the takeover phase into the adult stage, during which primary buds migrate to the colony center, open their siphons, and rapidly mature into adult zooids and becomes capable of independent feeding within just 36 hours (Sabbadin, Zaniolo and Majone, 1975). During takeover, adult zooids regress through apoptosis and phagocytosis, while emerging buds initiate differentiation and proliferation. Cellular debris is cleared by circulating phagocytes and reutilized by developing buds as part of a colony-wide recycling mechanism (Lauzon, Ishizuka and Weissman, 1992). Stem cell migration also occurs at this stage. Although somatic tissues experience weekly waves of apoptosis and phagocytosis, *B. schlosseri* colonies are capable of long-term regeneration and can live for years. This longevity is supported by the repeated trafficking of stem cells into new niches, which protects them from destruction and enables sustained self-renewal (Voskoboynik and Weissman, 2015). These dynamics suggest that the timing of cell cycle re-entry and tissue-specific differentiation is tightly regulated throughout the cycle particularly at the takeover stage.

### Functional Enrichment of Proliferation and Degradation Programs

Functional enrichment analysis of the two co-expression modules revealed their association with proliferation and degradation biological processes. Enrichment of DNA replication and cell cycle–related functions in the proliferation module supports the presence of tightly controlled mitotic programs in emerging buds (Ballarin and Manni, 2009). These processes are critical for maintaining genomic integrity and coordinating precise cell division across the colony. High telomerase activity reported in early budding stages further underscores the need for robust proliferative capacity to sustain continuous regeneration (Laird and Weissman, 2004). The proteasome also emerged as a central player, enriched in the proliferation module due to its role in protein turnover and cell cycle progression. This is consistent with its established function in regulating cyclins and cyclin-dependent kinase inhibitors via the ubiquitin-proteasome system (UPS) (Glickman and Ciechanover, 2002; Goldberg, 2003). In a colonial invertebrate like *B. schlosseri*, synchronized regression and renewal likely depend on such precise proteostasis. Supporting this, stress response and protein quality control pathways were co-enriched, suggesting that proteasome-mediated degradation maintains developmental fidelity under fluctuating physiological conditions (Tomanek, 2011). Enrichment of nucleocytoplasmic transport pathways further suggests active trafficking of regulatory proteins and RNAs between the nucleus and cytoplasm during rapid transitions. Ribosome biogenesis was also among the enriched processes during proliferative stages. Disruptions in ribosome production are known to impair cell growth and trigger cell cycle arrest, underscoring its importance in maintaining proliferative potential (Destefanis, Manara and Bellosta, 2020). As a tightly regulated and energy-intensive process, ribosome biogenesis supports sustained protein synthesis, which in turn fuels biomass accumulation, cell cycle progression, and differentiation. In proliferating cells, particularly during development or regeneration, upregulation of ribosomal RNA transcription, processing, and ribosomal protein production ensures sufficient translational capacity to meet the demands of rapid cell division (Thomas, 2000). These functional signatures in *B. schlosseri* reinforce the importance of translational control as a key node linking growth signals to cell cycle regulation and tissue homeostasis. Together, these enrichments point to coordinated upregulation of cell cycle progression, protein synthesis, and degradation machinery during takeover, all of which are essential for zooid renewal.

In contrast to the proliferation module, enrichment of ATP-binding cassette (ABC) transporters in the degradation module suggests increased membrane transport activity during zooid regression. These transporters may facilitate the removal of metabolic byproducts or the uptake of recycled nutrients, processes that are particularly important during large-scale tissue breakdown (Rees, Johnson and Lewinson, 2009). Additional enrichment of peroxisome activity, calcium signaling, and protein digestion and absorption pathways reflects the early activation of catabolic processes that enable the recycling of cellular components to support new tissue development (Ballarin, Schiavon and Manni, 2010). Peroxisomal proteins, which are involved in lipid β-oxidation and detoxification of reactive oxygen species, may help mitigate oxidative damage during tissue resorption (Schrader and Fahimi, 2006). This is especially relevant in *B. schlosseri*, where synchronous degeneration in densely packed zooids may create localized oxidative stress. Calcium signaling, which regulates apoptosis, mitochondrial dynamics, and cytoskeletal remodeling (Berridge, Bootman and Roderick, 2003), was also enriched during early regression stages but gradually declined. This trend may reflect the resolution of early remodeling events as regenerating tissues stabilize. Colonial ascidians often depend on localized recycling to sustain growth in nutrient-limited environments (Kürn *et al*., 2011). Accordingly, the observed decline in proteases and related enzymes involved in protein degradation over time likely represents a transition from catabolic resource mobilization to anabolic biosynthesis. The enrichment of catabolic and metabolic remodeling pathways in the degradation module, which is highly expressed in regressing zooids, is consistent with a prior apoptosis assay that reported a markedly higher rate of apoptotic cells in collapsing zooids, particularly within the gut epithelium and the pyloric gland (Tiozzo *et al*., 2006). The Functional modulations revealed by proteomics are consistent with a prior transcriptomic analysis that identified dynamic regulation of apoptosis-related genes across the blastogenic cycle, particularly during pre-takeover and takeover stages (Campagna *et al*., 2016). Key regulators such as caspase-2, AIF, and IAP were transcriptionally modulated in coordination with bud development and zooid regression. The current proteomic results extend these findings to the protein level, revealing parallel regulation of pathways involved in apoptosis, proteolysis, and cytoskeletal remodeling. These findings support the interpretation that zooid resorption is a tightly regulated, energy-coupled process involving coordinated degradation, recycling, and detoxification.

### Candidate Cell Cycle Regulators

Efficient cell cycle re-entry, particularly through the G1/S transition, is a major bottleneck in establishing proliferative primary cultures (Hume, Dianov and Ramadan, 2020). In this context, candidate *B. schlosseri* - specific cell cycle regulators that may support proliferation and extend the longevity of primary cultures were examined.

CDK1 is a known master regulator of the cell cycle, essential for driving cells through the G2/M transition and mitosis. It forms a complex with cyclin B to initiate mitotic entry by phosphorylating key substrates involved in chromatin condensation, nuclear envelope breakdown, and spindle assembly (Enserink and Kolodner, 2010). Beyond its core mitotic functions, CDK1 also phosphorylates p53 at Ser315, modulating its stability and transcriptional activity (Nantajit *et al*., 2010). These findings position CDK1 at a critical intersection between cell cycle progression and DNA damage response pathways. Moreover, CDK1 is also the sole CDK required for cell cycle progression in yeast (Nasmyth, 1993), and has been shown to drive the mammalian cell cycle in the absence of other CDKs (Santamaría *et al*., 2007). In this study, CDK1 displayed the most prominent stage-specific change in abundance, underscoring its potentially dominant role in orchestrating cell cycle progression during blastogenesis. Additional CDKs, including CDK5 and CDK20, were detected in this study, but did not show significant protein abundance changes. Interestingly, CDK4, a common target in mammalian cell cycle manipulation, was not detected in the study. This aligns with recent reports of CDKN2 (an inhibitor specific to CDK4/6) being absent in urochordates (Yuki *et al*., 2024), suggesting that the CDK4/6–CDKN2 regulatory axis may be reduced or absent in *B. schlosseri*. These findings support a model in which CDK1 plays a particularly central role in cell cycle regulation in this species, possibly compensating for the absence of canonical G1-phase regulators found in vertebrates, like CDK4. Together, CDK1 stands out as a prime candidate for functional manipulation in *B. schlosseri* cell cultures.

CDK2 is also one of the key regulators of the cell cycle, primarily functioning at the G1/S transition and during S phase progression. When bound to cyclin E or cyclin A, CDK2 drives DNA replication by phosphorylating targets involved in replication origin licensing and nucleotide biosynthesis (Honda *et al*., 2005). Although CDK2 is not strictly essential in some mammalian systems due to functional redundancy with CDK1, it plays a crucial role in fine-tuning S-phase entry and maintaining genome stability (Fagundes and Teixeira, 2021). In the context of *Botryllus schlosseri*, CDK2 is significantly upregulated during the TOB compared to TOZ, suggesting that CDK2 contributes to the proliferative capacity required for asexual reproduction. As such, CDK2 represents a promising candidate for functional manipulation to promote cell cycle re-entry in *B. schlosseri* primary cell cultures.

PCNA is among the most differentially expressed proteins observed in this study, highlighting its central role in the proliferative phase of the blastogenic cycle. As a DNA polymerase processivity factor, PCNA functions as a sliding clamp, tethering DNA polymerases to the template strand and ensuring efficient replication (Moldovan, Pfander and Jentsch, 2007). It is widely recognized as a conserved marker of active proliferation across eukaryotes (Iatropoulos and Williams, 1996). In addition to its essential function in DNA synthesis, PCNA also serves as a regulatory hub for several DNA repair pathways, including mismatch repair and translesion synthesis, thereby maintaining genome integrity during rapid cell division (Kelman, 1997). Its strong upregulation in TOB likely reflects both heightened proliferative activity and increased demand for genome maintenance. Given these roles, PCNA may serve not only as a mechanistic contributor to replication and repair but also as a valuable diagnostic marker for monitoring the success of culture conditions or genetic interventions aimed at promoting cell cycle re-entry and establishing long-term proliferative cell lines in *B. schlosseri*.

HDAC2 was significantly upregulated in TOB, suggesting a role in chromatin-mediated cell cycle regulation. As a class I histone deacetylase, HDAC2 removes acetyl groups from histones, promoting chromatin condensation and repression of cell cycle inhibitor genes. Loss of HDAC2 leads to G arrest and increased expression of inhibitors such as p21^CIP1^ and p57^Kip2^, demonstrating its importance in G /S progression (Segré and Chiocca, 2011). In addition to chromatin remodeling, HDAC2 also deacetylates non-histone proteins, implicating it in broader roles including DNA replication and repair (Miller *et al*., 2010). In *B. schlosseri*, its upregulation in primary buds likely reflects the heightened need for transcriptional control and genome maintenance during rapid cell proliferation. HDAC2 has also been extensively investigated for its role in cancer, where its overexpression is associated with tumor progression, poor prognosis, and resistance to therapy in multiple cancer types, including colorectal, gastric, and lung cancers (Weichert *et al*., 2008; Jung *et al*., 2012). While HDAC1 and HDAC2 often function redundantly in other systems, only HDAC2 was detected in this study, suggesting a potentially non-redundant and dominant role in regulating cell cycle transitions during bud formation. These characteristics position HDAC2 as a promising target for promoting cell cycle re-entry and supporting proliferation in primary cell culture systems.

BUB3 is a conserved spindle assembly checkpoint protein known for its critical role in monitoring chromosome alignment and inhibiting anaphase onset until all kinetochores are properly attached to spindle microtubules, thereby preserving genomic integrity (Larsen *et al*., 2007). It forms complexes with BUB1 and BUBR1 to ensure accurate mitotic progression. In *B. schlosseri*, BUB3 was upregulated in TOB, likely reflecting the increased need for stringent checkpoint control during rapid mitosis. Beyond its canonical mitotic role, BUB3 also contributes to genome stability during interphase by promoting efficient telomere DNA replication in complex with BUB1, preventing replication stress and telomere fragility (Li *et al*., 2018). This dual function highlights BUB3’s importance in both mitosis and S-phase regulation. Given that chromosomal instability is a major barrier to long-term culture viability, maintaining BUB3 activity in primary cultures may help support genomic fidelity, telomere integrity, and sustained proliferative capacity necessary for successful cell line establishment.

SKP1 was significantly upregulated in TOB and is a core component of the SCF (SKP1–Cullin–F-box) E3 ubiquitin ligase complex. This complex plays a central role in cell cycle regulation by targeting key regulatory proteins for proteasomal degradation (Xie, Wei and Sun, 2013). In yeast and mammals, SKP1 links F-box proteins to the ubiquitination machinery to promote degradation of cyclin-dependent kinase inhibitors such as Sic1 and p27, thereby facilitating G /S progression (Bai *et al*., 1996). Dysregulation of SKP1 and its associated SCF complexes has been observed in multiple cancers, including lung and prostate cancer. Recent studies have shown that pharmacological disruption of SKP1, can impair tumor growth and exhibits promising anti-tumor activity in preclinical models, highlighting the therapeutic potential (Li *et al*., 2025). Beyond cell cycle control, SCF complexes also mediate degradation of damaged or excess DNA licensing factors, contributing to genome stability and proteostasis (Silverman, Skaar and Pagano, 2012). In *B. schlosseri*, the upregulation of SKP1 may reflect a conserved regulatory axis that ensures rapid and orderly cell proliferation during blastogenesis.

SMC3, a core component of the cohesin complex, is essential for establishing sister chromatid cohesion during S phase and ensuring accurate chromosome segregation during mitosis. SMC3 undergoes a tightly regulated acetylation-deacetylation cycle at its nucleotide-binding domain, mediated by ESCO1/Eco1, which is critical for proper cohesion establishment and maintenance until anaphase(Beckouët *et al*., 2010). Beyond its canonical role in chromatid cohesion, the cohesin complex also participates in DNA repair and transcriptional regulation, linking chromatin architecture to gene expression (Wu and Yu, 2012). In *B. schlosseri*, elevated SMC3 expression during TOB likely reflects the increased demand for mitotic fidelity and genome stability during rapid proliferation.

Autophagy and Beclin 1 regulator 1 (AMBRA1), a known positive regulator of autophagy, was previously characterized in *B. schlosseri* as playing a role in tissue remodeling during the blastogenic cycle (Gasparini *et al*., 2016). AMBRA1 was also detected in the current proteomic study, although not significantly modulated between TOB and TOZ ((log FC =-0.96; FDR = 0.01). This modest change is consistent with the known regulatory mechanism of AMBRA1 as an intrinsically disordered scaffold protein, whose activity is predominantly controlled through post-translational modifications rather than large shifts in protein abundance (Cianfanelli, Nazio and Cecconi, 2015). AMBRA1’s known role in regulating both autophagy and cell proliferation in other systems (Maria Fimia *et al*., 2007; Maiani *et al*., 2021) suggests it may act as a context-dependent modulator during tissue remodeling in *B. schlosseri*.

Additionally, 14-3-3 proteins, which modulate the localization and activity of CDKs, phosphatases, and apoptosis regulators, may help coordinate cell cycle transitions during rapid proliferation (Mhawech, 2005). Their upregulation may help coordinate signaling during rapid cell cycle transitions. Although most of these proteins have been extensively studied in vertebrates, their conservation and regulation in *B. schlosseri* suggest functional similarity in checkpoint regulation and signal integration.

Finally, the present proteomic study identified significant upregulation of proteins involved in ribosome biogenesis and translation elongation in TOB compared to TOZ. These include elongation factors such as EEF2K, EEF1G, EEF2, EEF1D, and EEF1B2, suggesting a broader activation of the translational machinery to support proliferative demands. Activation of biosynthetic pathways is a hallmark of proliferative signaling, and in metazoans is commonly coordinated by the mTOR pathway, which directly enhances translation initiation and elongation through phosphorylation of factors such as S6K1 and 4E-BP1 (Hay and Sonenberg, 2004). Although RPS6KB2 (encoding S6K2) was detected in the study but not significantly differentially expressed, the broad upregulation of translation initiation factors such as multiple eIF3 subunits and elongation factors strongly support enhanced biosynthetic output consistent with mTOR pathway activity.

In parallel with mTOR-driven biosynthesis, other proliferative signaling networks may also contribute to enhanced translational output. Additionally, proliferative networks such as Wnt/β-catenin/Myc signaling amplify biosynthetic capacity by promoting ribosome biogenesis and global protein synthesis (He *et al*., 1998). Components of the Wnt/β-catenin pathway were not directly detected in the present study, likely due to their membrane localization or post-translational regulation. However, activated Wnt/β-catenin signaling was indicated by the CK1α downregulation observed in the growing bud (TOB) relative to the regressing zooid (TOZ). CK1α is a primary negative regulator of the Wnt/β-catenin axis as it phosphorylates β-catenin to trigger its degradation and is inhibited by Wnt/β-catenin activation in other models (Shen *et al*., 2025).

Together, these findings nominate CDK1, CDK2, HDAC2, and PCNA as key regulators of proliferation and promising candidates for functional manipulation. These proteins represent potential entry points for developing molecular tools to promote cell cycle re-entry, monitor proliferative capacity, and support the long-term establishment of cell cultures in colonial tunicates.

## Conclusion

This study provides the first proteome-wide map of the *B. schlosseri* blastogenic cycle at the individual animal level, revealing coordinated, stage-specific shifts in protein abundance that reflect synchronized tissue proliferation, differentiation, and regression. These data illuminate the molecular architecture of asexual development in a colonial chordate, identifying core cell cycle regulators and biosynthetic machinery that support bud formation and growth. Conversely, enrichment of senescence-and apoptosis-related proteins in regressing zooids highlights the developmental precision of programmed tissue removal. By linking proteomic dynamics to morphogenetic phases of the blastogenic cycle, this work establishes *B. schlosseri* as a powerful model for studying the balance between regeneration and degeneration in a naturally cycling developmental system.

Beyond these biological insights, this work lays a foundation for manipulating primary cells toward immortalization. The strong and specific expression of proliferation-associated proteins nominates candidates for functional testing, while PCNA emerges as a reliable molecular marker to track proliferation in vitro. Ultimately, this approach may enable the generation of the first immortalized cell line in a colonial tunicate, creating new opportunities for both regenerative biology and experimental developmental systems research. While this analysis focused on steady-state protein abundance, future studies incorporating post-translational modifications, such as phosphorylation and ubiquitination, will be essential to resolve dynamic regulatory mechanisms driving tissue remodeling and stem cell activity in this model.

## Materials and Methods

### Animal Husbandry

Wild colonies of *Botryllus schlosseri* were collected from floating docks at Berkeley Marina, California (United States). Larvae of these wild colonies were generated via sexual reproduction and attached as oozooids to glass slides at the UC Davis Cole B facility within one week after field collection. These lab-born colonies were then reared adhering to glass and kept vertically in 2.8 L glass tanks with 30ppt standing artificial sea water (ASW) at a constant temperature of 20 °C and aerated by air stones as described by Rinkevich and Shapira (Rinkevich and Shapira, 1998). All genotypes used in this study were raised from individually spawned, sexually reproduced oozooids and reared in separate tanks to avoid competition (Taketa *et al*., 2015). Colonies were fed twice a week with a combination of live algae (Dunaliella, Tetraselmis, Isochrysis and Nannochloropsis) and Roti-Rich Liquid Invertebrate Food (Florida Aqua Farms). ASW for each tank was fully changed every week, and colonies were gently cleaned once a week using soft brushes. All experimental colonies have been born and reared in stable lab conditions for at least 3 months, were in good health, and reproduced asexually via regular one-week blastogenic cycles.

### Dissection

Colonies were carefully cleaned and photographed before dissection under a stereomicroscope (Leica EZ4 W). To determine the appropriate stages, the development of buds and zooids was monitored daily. A healthy colony completes a full blastogenic cycle every 7 to 8 days. Using two sterile size 0 insect pins, the colony tunic was sliced open from the common atrial siphon to the oral siphon. Individual zooids were therefore exposed and carefully removed from the colony. Samples were kept on ice during dissection and snap-frozen in liquid nitrogen immediately afterwards, then transferred to a-80°C freezer for storage. A total of at least 50 zooids collected from each of the four blastogenic stages (A, B, C, and Takeover) and 70 primary buds from the takeover stage for each genotype were pooled together to ensure sufficient protein recovery. A total of three genotypes were used for the first study for five blastogenic stages and a total of seven genotypes were used for the second study for TOB, SAZ, and TOZ.

### Sample preparation

Sample preparations were performed as previously described (Kültz *et al*., 2024) with few modifications. Briefly, tissues were homogenized in lysis buffer (8M urea, 50mM Ambic) using 1mm zirconium beads (Benchmark D1032010) shaking in microtube homogenizer (Benchmark beadbag) at 3500 rpm for 30 seconds. Proteins taken from the supernatant were then reduced using 5mM dithiothreitol (DTT) for 10 min at 60C, alkylated with 15mM iodoacetamide (IAA) in the dark for 30 min. Remaining free IAA was quenched by further increasing DTT to 10mM. After protein quantification, urea was diluted with Ambic and subjected to trypsin digestion at a 1:50 trypsin:protein ratio at 37°C for 3 hours. Post-digestion, peptide cleanup was performed using Pierce C-18 spin columns (Thermo Scientific 89870) according to the manufacturer protocol.

Peptide concentration was quantified using Pierce fluorometric Quantitative Peptide Assay (Thermo Scientific 23290) before MS analysis.

### LC-MS/MS Acquisition

Liquid chromatography-mass spectrometry (LC-MS) acquisition were performed following established protocols for quantitative label-free proteomics (Kültz *et al*., 2024). Briefly, 100 ng of total peptide per sample was injected using a Bruker nanoElute® 2 UPLC system equipped operated in single column mode with a 25 cm x 150 µm x 1.5 µm Pepsep XTREME C18 reversed-phase analytical column (Bruker Daltonics 1893476). Peptide separation was achieved using a linear gradient of 3% to 33% acetonitrile in 0.1% formic acid over 60 minutes at a flow rate of 600 nL/min. The column temperature was maintained at 50°C. Mass spectrometry was performed using a UHD-QTOF mass spectrometer operating in positive ion mode (Bruker Impact II) interfaced online with the UPLC via a captive spray ionization source (Bruker CSI). For the first study, each sample was acquired twice, first in data-dependent acquisition (DDA) mode, and again using data-independent acquisition (DIA). For the second study, only DIA was used. For the first study, scan cycles for 74 scan windows (390 – 1130 m/z) were conducted at 10 m/z width ± 0.5 m/z overlap at a frequency of 50 Hz. For the second study, the same scan cycle parameters were applied except that each 1.5 sec scan cycle was preceded by acquisition of an MS1 spectrum obtained at the same scan rate (50 Hz) to enable annotation of corresponding precursor peptides and generation of spectral libraries with FragPipe (Kong *et al*., 2017) without the need for a separate DDA run.

### Data processing

Mass spectrometry raw DDA data for study 1 were processed using FragPipe 22.0, which includes the MSFragger search engine for peptide identification and quantification (Demichev *et al*., 2022). A spectral library was constructed from DDA runs and subsequently applied to DIA data of the same samples using Skyline (Pino *et al*., 2017) to extract and normalize (by sample median abundance) all transition peak areas. Spectral library filtering to remove interferences and non-diagnostic ions was performed as previously described (Kültz *et al*., 2024). For study 2, FragPipe 22.0 was used to generate a spectral library that was then imported to Skyline. For spectral library annotation, the predicted *B. schlosseri* reference proteome was based on a recently published update of this species’ genome (Thier *et al*., 2024).

### Statistical Analysis

For relative quantitation all transition peak abundances were exported from Skyline and then normalized and extrapolated to protein abundances using DirectLFQ (Ammar *et al*., 2023). Statistical comparisons were performed using the ProLFQua (Wolski *et al*., 2023) package in R, applying ANOVA with multiple testing correction (FDR < 0.1) for study 1, and ANOVA and linear models with empirical Bayes moderation for study 2 to improve detection of differentially abundant proteins. To assess overall proteomic variation, Principal Component Analysis (PCA) and hierarchical clustering were conducted on DAPs to visualize sample grouping and identify major sources of variation across conditions, using the R packages “mixOmics” and ‘edgeR’, respectively.

### Weighted Gene Co-expression Network Analysis

Co-expression patterns were examined using Weighted Gene Co-expression Network Analysis (WGCNA) (Langfelder and Horvath, 2008) to identify protein modules with coordinated expression changes across the blastogenic cycle. DAPs identified by ANOVA testing (FDR < 0.1) were selected for analysis after multiple testing correction. The soft-thresholding power (β) was determined based on the scale-free topology criterion, and the network was constructed using a “signed” network type, which considers only positive correlations between proteins. The Topological Overlap Matrix (TOM) was calculated with a signed TOMType, and the minimum module size was set to 20. Module detection was performed with the blockwiseModules function in R, and a merge cut height of 0.25 was applied to merge similar modules. Module-trait relationships were examined by correlating module eigengenes (MEs) with sample traits. To visualize the temporal dynamics of module expression across blastogenic stages, the average Log-transformed protein intensity for each module was plotted by condition. For each module, the mean Log intensity of all proteins assigned to that module was calculated per biological replicate, and the group average and standard deviation were plotted across stages.

### Functional Analysis

Protein sequences derived from the *Botryllus schlosseri* proteome were functionally annotated using a two-step approach. First, BLASTP searches were performed against the SwissProt database using DIAMOND, with no taxonomic restrictions to assign functional annotations. Only annotations with an E-value < 10^-3^ were considered. In addition, eggNOG (Jensen *et al*., 2008) was used to identify orthologous relationships and retrieve KEGG (Kyoto Encyclopedia of Genes and Genomes) pathway annotations.

Functional enrichment analyses were performed using the clusterProfiler (Xu *et al*., 2024) R package to identify over-represented KEGG pathways within each co-expression module identified by WGCNA. Over-representation analyses were conducted using a customized functional database derived from eggNOG annotations. The background set included all proteins from the first study, and protein sets analyzed corresponded to those within each co-expression module identified by WGCNA. KEGG pathway enrichment was assessed using the *enrichKEGG* function, with the organism parameter set to “ko” (KEGG orthology). Statistical significance was assessed using the Benjamini-Hochberg (BH) multiple testing correction method, with pathways considered significantly enriched if the adjusted p-value (FDR) was < 0.1.

To reduce redundancy in enriched pathways, affinity propagation clustering was performed on the significant pathways using the R package “APCluster” (Bodenhofer, Kothmeier and Hochreiter, 2011). This method grouped similar pathways based on shared protein sets, retaining only the most representative ones. Enriched pathways were visualized using R package ggplot2. Pathways that were vertebrate-specific, human disease-specific, or biologically irrelevant to *B. schlosseri* were manually filtered out prior to visualization.

## Supplementary methods

### Genotyping

To confirm that the biological replicates used in this study represented independent genotypes, a panel of 12 polymorphic loci, the fusion-histocompatibility, or fuhc, was tested across all seven *B. schlosseri* colonies. The fuhc loci is responsible for allorecognition between colonies and encompasses the highly variable fester gene family, with a range of 7-13 alleles per individual according to recent study (Rodriguez-Valbuena *et al*., 2024 preprint). Here, each colony was found to have 4 alleles on average at different combinations of loci, confirming each colony represents a distinct genotype. Colonies were bred from wild populations and maintained separately under laboratory conditions to ensure consistent environmental exposure during the experimental timeline. Genomic DNA was extracted from tissue samples from each colony using the PureLink Genomic DNA Mini Kit (ThermoFisher Scientific) following the manufacturer’s protocol with a modified 7-hour incubation period. PCR was performed in 25 μL reaction volumes containing 12.5 μL of 2x EmeraldAmp Max HS PCR Master Mix (Takara), 0.5 μL each of forward and reverse primers for each locus, and 11.5 μL of DNA sample with DI water (sample volumes were calculated depending on individual DNA concentration to yield 50 ng of DNA and supplemented with water to reach 11.5 μL). The thermal cycling protocol included an initial denaturation step at 94°C for 3 minutes, followed by denaturation at 98°C for 10 seconds, annealing at locus-specific temperatures for 30 seconds, extension at 72°C for 30 seconds, a 30 cycle repeat and a final extension at 72°C for 5 minutes. PCR products were visualized using gel electrophoresis on a 1.5% agarose gel stained with SYBR Safe DNA Gel Stain (ThermoFisher Scientific) and imaged using UVP ChemStudio PLUS (Analytik Jena) with the SYBR Safe emission filter. Allele sizes were compared against the GeneRuler 50 bp DNA Ladder (ThermoFisher Scientific) to identify polymorphic patterns. The presence of unique banding patterns at multiple loci, as seen in representative gel images included in Supplemental Fig. S2, confirms the presence of distinct genotypes.

## Acknowledgement

We thank Baruch Rinkevich (Israel Oceanography & Limnological Research, National Institute of Oceanography) and Ayelet Voskoboynik (Hopkins Marine Station, Stanford University) for the advice on rearing *Botryllus schlosseri* and dissection. We also thank Stefano Tiozzo for the improved reference proteome.

## Data Availability

All MS proteomics data and metadata have been deposited and are publicly available in PanoramaPublic (https://panoramaweb.org/vwd01kl.url) and ProteomeXchange (PXD065460).

## Supplemental Figure Legends

**Supplemental Figure 1. Phylogenetic placement of *Botryllus schlosseri* among model organisms.** A phylogenetic tree illustrates the evolutionary relationships between major model organisms, highlighting the position of *B. schlosseri* within the chordate phylum. As a basal chordate diverging approximately 535 million years ago, *B. schlosseri* provides a valuable system for studying the evolution of regenerative capacity, development, and cell biology in relation to vertebrates. Created in BioRender. Dong, V. (2025) https://BioRender.com/c4zu2is.

**Supplemental Figure 2. Genotypic differentiation of *Botryllus schlosseri* colonies using 12 *fester* loci.** PCR results are shown for 12 *fester* gene primer sets across seven *B. schlosseri* genotypes. Each column represents a distinct *fester* locus, and each row corresponds to a unique genotype. Genotypic identity is determined based on the unique combination of presence or absence patterns across the loci, enabling discrimination between colonies.

