## Supplementary figures and images for "Proteome Dynamics Across the Blastogenic Cycle of *Botryllus schlosseri* Reveals Targets for Cell Immortalization"

### Supplemental Figure S1

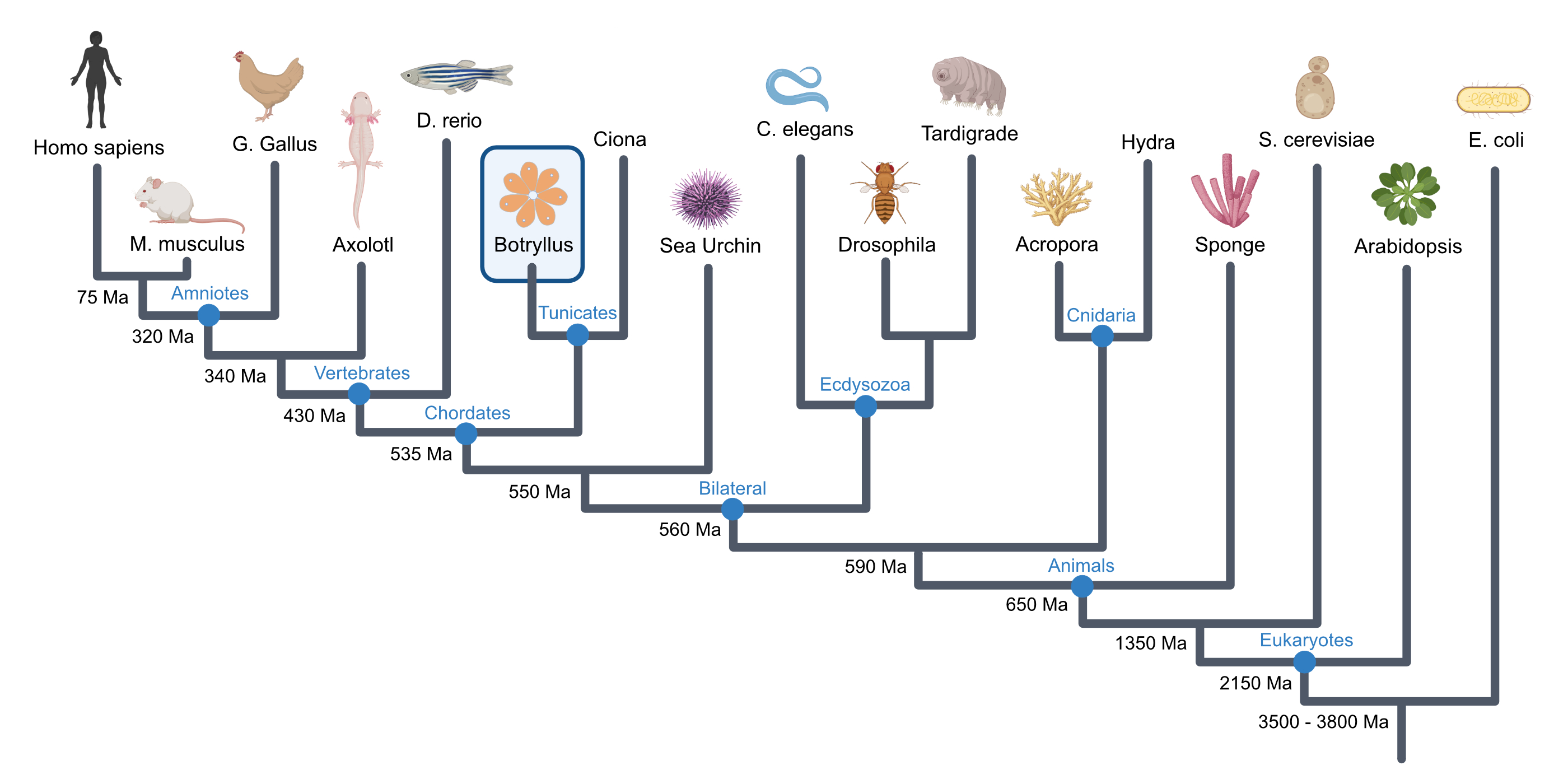

### Supplemental Figure S2

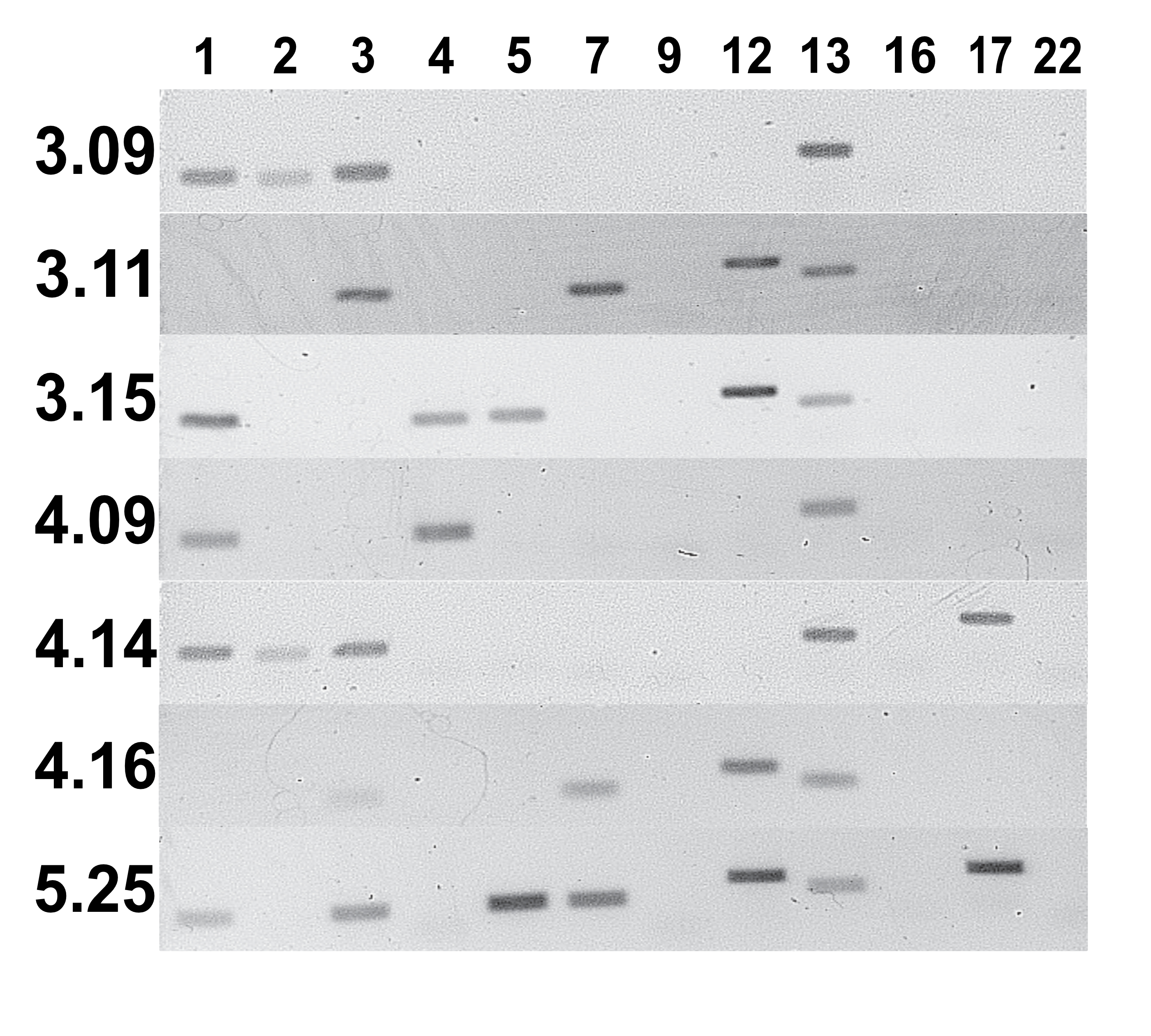
